# Coupling magnetically induced electric fields to neurons: longitudinal and transverse activation

**DOI:** 10.1101/275073

**Authors:** B. Wang, W. M. Grill, A. V. Peterchev

## Abstract

We present a theory and computational models to couple the electric field induced by magnetic stimulation to neuronal membranes. The response of neuronal membranes to induced electric fields is examined under different time scales, and the characteristics of the primary and secondary electric fields from electromagnetic induction and charge accumulation on conductivity boundaries, respectively, are analyzed. Based on the field characteristics and decoupling of the longitudinal and transverse field components along the neural cable, quasi-potentials are a simple and accurate approximation for coupling of magnetically induced electric fields to neurons and a modified cable equation provides theoretical consistency for magnetic stimulation. The conventional and modified cable equations are used to simulate magnetic stimulation of long peripheral nerves by circular and figure-8 coils. Activation thresholds are obtained over a range of lateral and vertical coil positions for two nonlinear membrane models representing unmyelinated and myelinated axons and also for undulating myelinated axons. For unmyelinated straight axons, the thresholds obtained with the modified cable equation are significantly lower due to transverse polarization, and the spatial distributions of thresholds as a function of coil position differ significantly from predictions by the activating function. For myelinated axons, the transverse field contributes negligibly to activation thresholds, whereas axonal undulation can increase or decrease thresholds depending on coil position. The analysis provides a rigorous theoretical foundation and implementation methods for the use of the cable equation to model neuronal response to magnetically induced electric fields. Experimentally observed stimulation with the electric fields perpendicular to the nerve trunk cannot be explained by transverse polarization alone and is likely due to nerve fiber undulation and other geometrical inhomogeneities.

## INTRODUCTION

### Background and motivation

Stimulation with a magnetically induced electric field (E-field) is a noninvasive technique that elicits or modulates neural activity. Transcranial magnetic stimulation (TMS) is widely used in neuroscience as a tool for probing brain function and connectivity (1). TMS is approved by the U.S.A. Food and Drug Administration (FDA) for the treatment of depression and migraine, as well as for pre-surgical cortical mapping, and is under study for other neurological and psychiatric disorders (2, 3). Magnetic stimulation of peripheral nerves, which is also FDA cleared, is used for nerve conduction testing, neuromodulation, and neurorehabilitation (4–7). The mechanisms determining the neural response to magnetic stimulation are still unclear, and experimental studies rely heavily on indirect, non-invasive measurements such as brain imaging and downstream neuromuscular responses (8, 9). Further, the strong electromagnetic coupling between the stimulus and electrophysiological recording systems presents challenges to direct *in vivo* recording from neurons and recording latencies are typically longer than ∼ 1 ms after the stimulus (10).

Computational models of neuronal activation by magnetically induced E-fields provide an approach to understand stimulation mechanisms and to optimize stimulation parameters (11–15). Like electrical stimulation, a two-stage approach is commonly used to simulate magnetic stimulation (16). In the first stage, the macroscopic E-field distribution is calculated under the quasi-static assumption (17, 18), for example by the finite element method (19, 20). The applied E-field 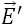 in magnetic stimulation includes a non-conservative primary source 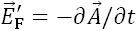 due to induction (21) and a secondary source 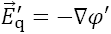 from charges associated with conductivity inhomogeneities (19, 22), which are represented by a vector potential 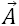 and a scalar potential *φ*′, respectively. In the second stage, the E-field is coupled to the cable equation (CE) that describes the neuronal response to stimulation (13, 14, 23, 24). The activating function *f*, proportional to the first spatial derivative of the E-field in the longitudinal direction of a neuronal process (typically an axon), is widely accepted as the term driving neural polarization (25). For magnetic stimulation, a hybrid CE was derived with the activating function generalized to include both the primary and secondary source E-fields explicitly (23, 24, 26)

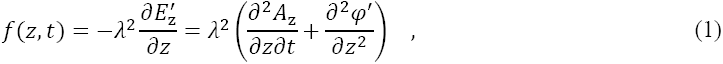

where *z* and λ are the axon’s longitudinal direction and length constant, respectively.

However, this approach raises several questions regarding the coupling between the two stages. Problems common to electrical and magnetic stimulation include the use of one-dimensional (1-D) cable representations of 3-D axons. While this simplifies the computational implementation, the interaction between the field and the membrane as well as some aspects of the membrane polarization are not captured. Also, the E-field in the first stage is obtained with macroscopic conductivity values and therefore may not reflect the detailed field distribution on cellular scales (27, 28). Issues specific to magnetic stimulation are introduced in the following two sections, and these require resolution to improve the rigor and utility of magnetic stimulation models.

### Coupling of E-field to neuronal membrane in magnetic stimulation

Although CEs are used for magnetic stimulation (23, 24, 29), their theoretical justification and computational implementation have limitations. First, the use of potentials in magnetic stimulation models should be reevaluated. The source scalar potentials *φ*′ can be ignored if the model has no external boundaries (29–31) or the primary source field is parallel to the boundary (23, 32–34). In other cases, however, *φ*′ is incorrectly ignored, for example using field distributions for peripheral nerves to study activation of cortical neurons (35). In response to the applied field, the cells generate a secondary E-field due to charge redistribution on the membrane, and this can be represented by a scalar potential,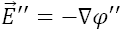 The response potential is typically neglected assuming its amplitude is small compared to the source field (29), and only the latter is included in the conventional 1-D CE for magnetic stimulation (24). The exclusion of the response potential is justified theoretically in the conventional CE because the membrane potential it describes is the mean value averaged around the circumference of the cable and not directly affected by the response field (36, 37). However, the response field needs to be included to describe the behavior of the neural cable in the transverse dimension and/or ephaptic interactions with neighboring membranes (27, 28, 36–40). The vector potential is used together with the scalar potential, especially the transmembrane potential, assuming that the latter behaves the same with a non-conservative E-field present (29). In the CE, the vector potential in the intracellular space is sometimes substituted with its extracellular value in analogy to the scalar potential (24). While these are valid numerical approximations, their theoretical rigor requires further evaluation.

Furthermore, a simple and accurate computational implementation of magnetic stimulation in CE solvers has not been established. The applied E-field, and especially the magnetically-induced component, is sometimes converted into an equivalent intracellular current injection (13, 41–43). The activating function at nodes of myelinated axons can be calculated using the integral of the E-field over neighboring internodes. However, these quantities are defined separately for each internode without establishing a global variable defined over the entire cell (32). As an alternative approach, Goodwin and Butson directly applied the total E-field solution obtained via the finite element method to cell models in the NEURON software, and hence implicitly defined a global E-field integral (14). However, no other details were provided for the method, such as its computational implementation or distinction from electric potentials.

### Activation of long nerves by transverse field in magnetic stimulation

Experiments with magnetic stimulation of peripheral nerves *in vivo* (44–47) and *in vitro* (48, 49) showed cases of neural activation that were inconsistent with predictions by the activating function. To explain this discrepancy, the transverse component of the E-field was proposed to contribute to neural activation (44, 50), and Ruohonen et al. introduced a modified activating function *f*_*m*_ (44) by adding the steady-state transverse polarization of cylindrical fibers (51, 52)

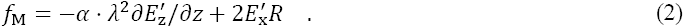

Here α is a scaling factor, *R* is the axon’s radius, and the x axis is aligned with the transverse direction of the E-field. The experimentally fitted α suggested an equal or even dominant contribution to activation by the transverse components of the field (44).

However, whether the transverse field component indeed activates peripheral axons is unknown, since the modified activating function (44, 53, 54) and other studies (30, 31) used simplifying assumptions including linear membrane models, unmyelinated axons, and neglecting the temporal integration required for membrane depolarization. We derived a modified CE (37) showing that modulation of the transmembrane potentials by the transverse field averages out to zero around the circumference of any neural compartment with a linear membrane (see Eqs. 6 and 7), thereby invalidating the use of the modified activating function as a simple predictor for neural activation. Moreover, our simulations with nonlinear membrane models showed that the effect of the transverse field on activation thresholds is much smaller than estimated by linear membrane models, especially for myelinated axons (37). Therefore, the modified CE should be used to examine whether transverse field components induced by magnetic stimulation affect activation thresholds and to what extent.

An alternate hypothesis to explain the experimental results is nerve undulation. Peripheral nerves exhibit undulation to accommodate compression and tension due to movement (55–57). The undulation of nerve fibers introduces short axonal segments that partially align with the E-field transverse to the nerve trunk, and could be the mechanism underlying so-called transverse-field activation (49, 54, 58). Previous studies of the effects of nerve undulation either used linear membrane models or investigated nonlinear membrane models for a limited number of coil types and positions. We examine activation due to undulation with that due to transverse fields in a straight axon under the same conditions, allowing an assessment of the relative contribution of the two activation mechanisms.

### Aims and organization of the paper

We present theoretical analyses and computational simulations to address the two questions described above, namely the coupling of magnetically-induced potentials to neuronal membranes and transverse-field activation.

The use of potentials is resolved by considering the electromagnetic–neuronal coupling on different time scales. The theoretical framework first examines charge storage on the neuronal membrane on the sub-ns time scale. Then, the neuronal response to the transverse field on the sub-μs time scale (52), i.e., transverse polarization, is described. The neural cable behavior on the μs to ms time scale is described by the CE. Specifically, the modified CE previously developed for electrical stimulation (37) is based on the asymptotic expansion of different temporal scales (36) and incorporates the fast response to the transverse E-field. The E-field coupling for magnetic stimulation is analyzed within the context of transverse polarization and the modified CE to provide more rigorous theoretical justification for the use of CE for magnetic stimulation (24). Moreover, quasi-potentials, a potential-like variable based on the integral of the E-field, are used in this theoretical analysis and for computational simulations of magnetic stimulation.

The transverse-field activation of peripheral nerves by magnetic stimulation is quantified using the modified CE and quasi-potentials. Unmyelinated and myelinated straight axons are simulated to obtain thresholds with both the conventional and modified CEs. The undulation of myelinated axons is also considered as an alternative factor contributing to activation by the transverse component of the E-field. The spatial distributions of the activation thresholds for circular and figure-8 coils for various positions with respect to an axon are presented, comparing modified versus conventional CEs for straight axons or undulating versus straight axons using the conventional CE.

## METHODS

### Theoretical framework of electromagnetic–neuronal coupling

#### Response of neuronal membrane on different time scales

The cell membrane is usually modeled as a distributed capacitor that separates charge and allows ionic current to pass through. For a linear membrane, the relationship between the reduced transmembrane potential *φ*_m_ and transmembrane current density *i*_m_ is described by

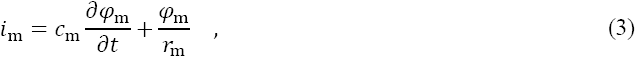

where *c*_m_ and r_m_ are the specific membrane capacitance and resistance, respectively. For a membrane at rest, *φ*_m_ can be considered a zero state response to the intra-and extra-cellular current densities normal to the membrane, which are considered to be continuous with *i*_m_ (current continuity, i.e., Eq. A1 in Appendix A). However, similar to boundaries of macroscopic conductivity changes on which surface charge density accumulates (19, 21, 22), the highly-resistive cell membrane is a physical boundary that can also store a net charge in response to applied E-fields and act as a source for a secondary E-field. The stored charge *q*_s_ evolves on sub-ns time scales and is instantaneous compared to the differential charge *q*_m_ accumulation involved in membrane polarization (see Appendix A). Charge storage generates an additional membrane polarization *φ*_m,zi_ (a zero input response) that is not part of Eq. 3. Although this additional polarization typically has negligible amplitude and does not affect computational models, its existence complicates the theoretical consideration in magnetic stimulation since the source field contains a non-conservative component. This issue is addressed in the next section.

When the cell responds to an applied E-field, the membrane differentially polarizes on the anodal and cathodal sides with a time constant on the order of 10 to100 ns (37, 38, 52), resulting in transverse polarization (51, 52, 59). For transverse polarization, the redistribution of the local membrane charges via the intra-and extra-cellular spaces does not change the mean *φ*_m_ of the whole cell, but shields the applied field in the intracellular space, resulting in a uniform intracellular potential *φ*_i_ that evolves on a slower time scale according to the cell’s electrophysiological state (37, 52). For a long cylindrical axon of radius *R*, the response to an extracellular E-field can be asymptotically decomposed (36) and the transverse polarization at any longitudinal location *z*is

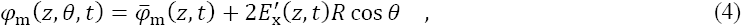

where the *x*-axis is locally defined by the direction of the transverse component of the quasi-uniform exogenous E-field and *θ* is the azimuthal angle on the axon’s circumference. The average transmembrane potential for any longitudinal location is 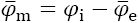 The mean extracellular potential 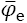 is specified to yield a unique solution, which is also termed the normalization condition (52), and is the same if calculated only for the primary source field without accounting for the presence of the cell 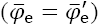 (37, 52). Along the axon,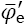 varies according to the longitudinal component of the exogenous field

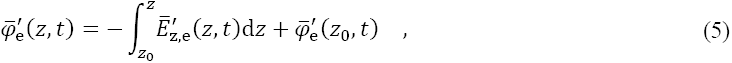

where the z-direction is the local longitudinal direction, *z*_0_ is the reference point on the axon, and 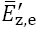 is the longitudinal component of the extracellular E-field averaged around the axon’s circumference (37).

The behavior of the neural cable on the μs to ms scale is described by the CE, for which the conventional form only includes the longitudinal dimension. Previously, we modified the CE to incorporate the transverse dimension into the 1-D longitudinal equation by assuming a local uniform-field solution (Eq. 4) for transverse polarization

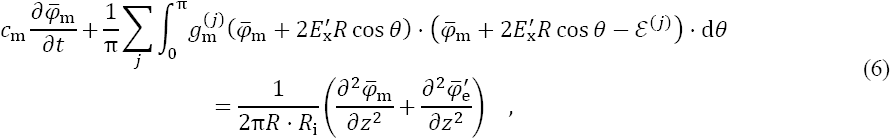

where *R*_i_ is the longitudinal resistance that relates the longitudinal current *I*_i_ and its driving intracellular E-field (*R*_i_*I*_i_ *= E*_z,i_), and the summation is over different types of ion channels *j*, each having reversal potential *ε*^*(j)*^ and nonlinear conductance 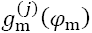 (37). By converting the *θ*-dependent ionic currents into an equivalent channel with parameters 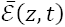 and 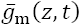 (37), the modified CE (Eq. 6) can be reduced to the form of the conventional CE

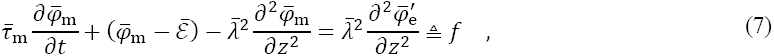

with location-and time-dependent length and time “constants”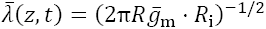 and 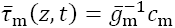. Since the extracellular potential varies around the neural cable, the activation by the longitudinal E-field should be calculated according to Eq. 5, using the primary extracellular field averaged around the neural compartment. The transverse-field activation is through the integration of ionic currents in Eq. 6 or equivalent parameters 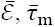 and 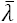 in Eq. 7. The contribution of the transverse E-field is complex for a nonlinear membrane, but averages out to zero for a linear membrane (37). Therefore, the simple addition of transverse and longitudinal E-field components in the modified activating function (Eq. 2) is invalid, whether linear or nonlinear membranes are considered.

#### Quasi-potentials and cable equation for magnetic stimulation

In the previous section, the analyses are presented using scalar potentials for electrical stimulation. However, the E-field induced by magnetic stimulation is non-conservative in nature and cannot be described by scalar potentials. The issue of using scalar potentials can be resolved, however, by exploiting the low spatial variation of the induced E-field on microscopic scales. Both the transverse and longitudinal components of 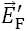 can be considered quasi-uniform at any location on the neural cable (60), and the transversely-uniform longitudinal component varies slowly along the cable’s axis. Therefore, *quasi-potentials ψ* can be defined along the neural cable to combine the effect of both 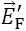 and 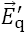

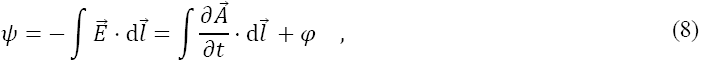

in which the line integral starts from a point in the intracellular space and the E-field includes both the source and response fields. In any transverse cross-section of the neuron, *ψ*_i_ is spatially uniform like *φ*_i_ due to the membrane’s response field canceling the source field (37), and *ψ*_m_ is the same as *φ*_m_ because the additional polarization component *φ*_m,zi_ is eliminated in this integration. Therefore, transmembrane quantities such as polarization and ionic current densities can still be described in their original forms (29). Similar to 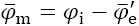 in the transverse polarization analysis, the average *ψ*_m_ relates the intra-and extra-cellular quasi-potentials

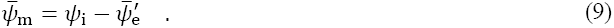

Therefore, only 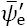 needs to be calculated at any longitudinal location and quasi-potentials can be considered an extracellular field parameter defined along the 1-D cable, with longitudinal variation given similar to Eq. 5

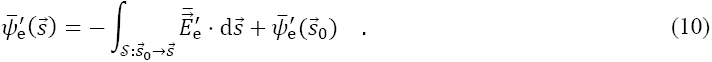

Here,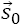 is the location of the reference point on the neural cable to start the integration (typically the soma or the (main) axonal terminal, for convenience),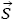 is the location of interest, 𝒮 is the path of the integral from 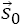 to 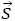 along the neuron’s topology (i.e., along the sequence of neuronal segments between 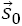 and 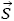, rather than a straight line between the two points through extracellular space), and d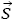 is the differential vector along 𝒮. As neurons typically have a tree-like topology that do not form connected loops on themselves, quasi-potentials are well-defined for individual neurons and should be calculated independently for each cell. Due to the low spatial variation of the E-field, the integration step is mostly determined by the morphology of the cell, e.g., distance between branching points and curvature of bends. For numerical computation, quasi-potentials can be approximated by traversing the cell’s topology with sufficiently high resolution and adding the source E-field component along each neural process in discrete steps

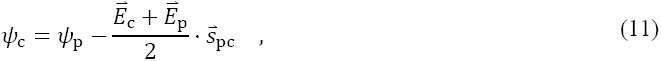

where the subscripts c and p indicate the current (child) compartment and its previous (parent) compartment in the tree topology of the discretized neuron model (with the root of the tree having *ψ*_0_ = 0); 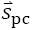 is the displacement vector between them; and the notations for source field, averaging, and extracellular domain are ignored for simplicity.

The hybrid CE for magnetic stimulation, which uses an activation function defined by Eq. 1, has the correct mathematical form (24) but its theoretical justifications has limitations, especially accounting for transverse polarization. The transverse primary source - ∂*A*_x_/∂*t* is omitted and thought not to contribute to transmembrane current (24), whereas its contribution is implicitly contained in the form of 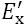 when transverse polarization and the modified CE are considered. The longitudinal primary source - ∂*A*_z_/∂*t* is an additional term driving the intracellular current within the cable (23, 24, 26)

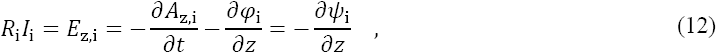

and the extracellular value *A*_z,e_ is simply substituted for *A*_z,i_, as they are considered identical due the small radial distance between the two domains (24). Although the variation of the field is indeed small, the rigorous interpretation of this substitution is given by Eqs. 9 and 10, which together account for the averaging of extracellular quasi-potentials around the cable’s cross-sectional circumference and the relationship of the average potentials in the longitudinal direction. Therefore, magnetic stimulation can use the CEs for electrical stimulation (Eqs. 6 and 7) by directly replacing the extracellular scalar potentials with extracellular quasi-potentials, which provides significant convenience over the equivalent hybrid CE or other computational methods (see Discussion).

### Computational quantification of transverse-field activation by magnetic stimulation

The simulations were performed using custom code in MATLAB (versions 2016a and 2017b, The Mathworks, Inc., Natick, MA, USA). The code and data are available online at https://doi.org/10.5281/zenodo.1186947.

#### Simulation setup for magnetic coils

We quantified magnetic stimulation of peripheral nerves with a transverse E-field using the modified CE and nonlinear membrane models. The coils were placed parallel to the interface of a semi-infinite volume conductor in which the nerve was positioned parallel to the interface (Fig. 1A). No secondary E-field was induced within the volume conductor and the analytical solution for idealized circular coils (61) was used to calculate the E-field for a given rate of change of the coil current. The circular coil had a 5 cm diameter and 21 co-localized turns, similar to the one used by Ruohonen et al. (44). The figure-8 coil was simulated by combining the field solutions of two circular windings each with a 4 cm diameter and 14 turns (44), in which the current directions were opposite, and the coil was orientated with the peak field either aligned with or perpendicular to the nerve. The E-field was modulated temporally with either monophasic or half-sine waveforms recorded from a MagPro X100 device (MagVenture A/S, Farum, Denmark; Fig. 1B). Both waveforms had peak amplitudes at pulse onset normalized to unity, a first phase with approximately 75 μs duration, and a time integral of zero. The choice of the half-sine waveform was due to its similarity to biphasic waveforms in electrical stimulation, whereas a magnetic “biphasic” waveform has three phases and a significantly longer duration (62).

**FIGURE 1.**
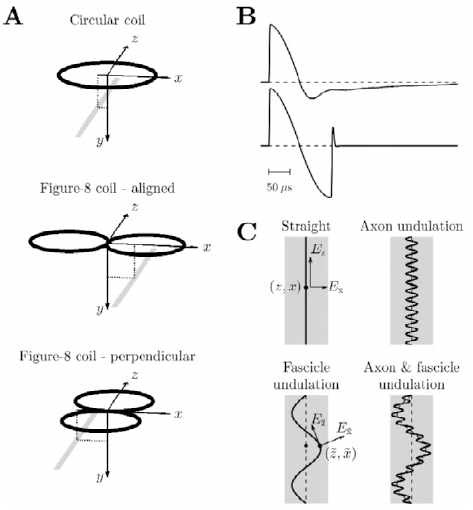
Simulation setup. **A**: Three configurations of magnetic stimulation coils (black) with a nerve (gray) underneath. The coordinate system shows the nerve’s distance to the coil center in the vertical (*y*) and lateral (*x*) directions. **B**: E-field waveforms of monophasic and half-sine magnetic stimulation pulses, with peak amplitude normalized to unity at pulse onset. **C**: Four types of axon placement within the nerve trunk. The wavelengths and amplitudes of the undulation are exaggerated for visualization.

#### Straight axons

The Hodgkin-Huxley (HH) model (63) and Richardson-McIntyre-Grill (RMG) model (64) were used to model unmyelinated and myelinated straight axons, respectively (Appendix B). The axon had a length of about 30 cm and its midpoint was aligned with the coil’s center in the *z*-direction (Fig. 1A). The axon’s distance to the coil was varied vertically (y) between 0.5 cm to 4 cm, and the lateral distance (x) for the three coil configurations was between −5 cm to 5 cm, −2.5 cm to 7.5 cm (range shifted due to symmetry of field distribution), or −5 cm to 5 cm, respectively, all with 2.5 mm intervals.

The E-field calculated along the axon (Fig. 1C) was coupled to our custom modified CE solver directly for the transverse component *E*_x_ (for myelinated axons, at Ranvier nodes only) (37) and using quasi-potentials for the longitudinal component *E*_z_. The simulation time step was 5 μs (HH model) or 2 μs (RMG model) and the membrane azimuthal discretization was set to 15 steps within *θ* ∈ [0, Π] for the modified CE. To avoid action potential initiation at the axon terminations, the activating function was set to zero at the ends of the axon. Activation thresholds were determined with 0.5% accuracy as the stimulation amplitude that elicited an action potential propagating to the axon terminal in the positive *z*-direction and reported in units of amperes per microsecond (A/μs) for the coil current at the pulse onset. To quantify the effect of transverse polarization on activation thresholds, the percent difference of thresholds between the modified CE and the conventional CE was calculated.

#### Undulating axons

We simulated undulation of nerve fibers with the myelinated RMG axons using the conventional CE. The unmyelinated axons were excluded because they have very small diameters in mammalian nerves and therefore thresholds too high to be excited by magnetic stimulation. The modified CE was not used due to the negligible influence of the transverse field on the thresholds of myelinated axons (see Results section and (37)). The positioning of the nerve trunk (x, y) was the same as for the straight axon, and the undulation was assumed to be sinusoidal within the x-z plane (Fig. 1C), consisting of a relatively short-wavelength undulation of the axon within the fascicle and a longer-wavelength undulation of the fascicle within the nerve trunk (54)

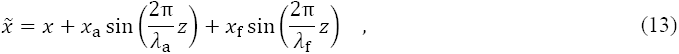

with *z*as the position along the nerve trunk, and *x*_a,f_ and *λ*_a,f_ the amplitudes and wavelengths of the undulations with subscripts a and f representing the axon and fascicle, respectively. The axon undulation had an amplitude of *x*_a_= 40 μm and a wavelength of *λ*_a_= 0.2 mm (49, 55, 57, 58). The E-field distribution along the axon was insensitive to a shift of the axon undulation because its wavelength was small compared to the spatial variation of the field, and therefore its phase was set to zero. The fascicle undulation had an amplitude of *x*_f_ = 0.8 mm and a wavelength of *λ*_f_ = 5 cm, and its phase was set to zero to maximize the amplification of the activating function (54). To investigate the contribution of the two undulation components, simulations were also performed on axons with either the axon or fascicle undulation only. For the undulating axons to have the same internodal distance as the straight axon, the arc length of Eq. 13 was first calculated as a function of *z*. Then, the inverse relationship between arc length and *z*was used to convert the compartment positions of a straight axon (x and *z*) to the corresponding undulating axon (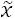 and 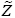). The longitudinal E-field 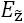 was calculated according to the position and local orientation of the undulating axon and converted to quasi-potentials. The discretization along the axon was increased by 10 times for the internodes to capture the increased spatial variation. To mitigate the increased polarization near the terminals due to the undulation, the axon’s length was extended by 20% in each direction without changing the location for action potential detection, and the surface area of the new terminal nodes was enlarged by a hundred times to further suppress terminal polarization. All other neuronal parameters were the same as the straight axons (Appendix B). The effect of the undulation was quantified as the percent difference of thresholds between the undulating and straight axon, both obtained with the conventional CE.

## RESULTS

### Distributions of electric field components relevant to neural activation

The distributions of the E-field calculated for the three coil configurations (Fig. 2), are in agreement with theoretical expectations and the results of Ruohonen et al. (44). The gradient of the longitudinal field and strength of the transverse field were largest for similar longitudinal locations along the nerve (*z* = ±2.5 cm for the circular coil, ±2 cm for the aligned figure-8 coil, and ±4 cm and 0 cm for the perpendicular figure-8 coil). In the lateral direction, the longitudinal field gradient was largest for the nerve trunk positioned tangential to the circular coil windings (*x* = ±2.5 cm for the circular coil, ±4 cm and 0 cm for the aligned figure-8 coil, and ±2 cm for the perpendicular figure-8 coil), whereas the transverse field’s amplitude was largest for the nerve trunk passing near the center of the windings (*x* = 0 cm for the circular coil and perpendicular figure-8 coil, and ±2 cm for the aligned figure-8 coil).

**FIGURE 2.**
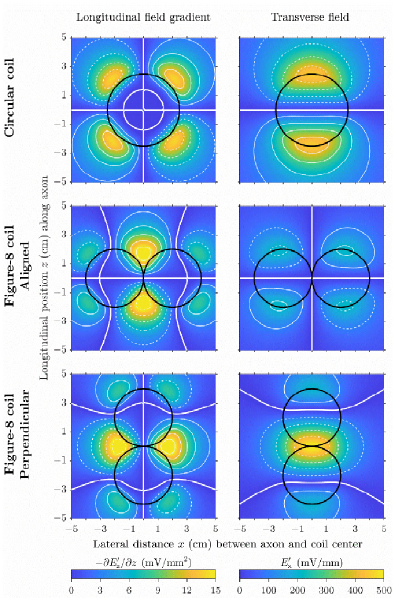
Distributions of E-field components contributing to neural activation in a plane 1 cm below a circular coil (top row) and a figure-8 coil aligned with or perpendicular to the nerve (middle and bottom rows, respectively). The coil current has a rate of change of 100 A/μs. The gradient of the longitudinal field (left column) and strength of the transverse field (right column) are shown along the nerve (*z*) for different lateral locations (x) relative to the coils (black outlines). The white contour lines, spaced at 3 mV/mm^2^ and 100 mV/mm intervals, respectively, show positive values with thin solid lines, zero with thick solid lines, and negative values with dashed lines.

### Transverse stimulation of unmyelinated straight axons

The thresholds for excitation of straight unmyelinated HH-model axons with monophasic and half-sine magnetic pulses were on the order of thousands of A/μs or more (Fig. 3), infeasible for practical magnetic stimulation devices. The conventional CE (left column) shows that thresholds were smallest at positions where the activating function was largest, i.e., at the edges for circular coils (top group) and perpendicularly orientated figure-8 coils (bottom group) as well as at the center for aligned figure-8 coils (middle group). The threshold increased with vertical distance from the coil and also when the axon was moved laterally. When aligned through the center of the coil windings, the activating function along the axon was zero for the circular coil or perpendicular figure-8 coil, and no action potential could be generated; for the aligned figure-8 coil, thresholds were much higher under the center of each wing.

**FIGURE 3.**
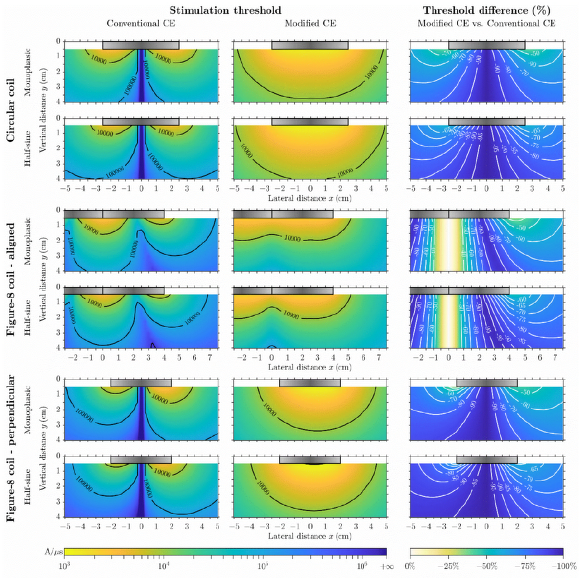
Activation thresholds of straight unmyelinated HH-model axon for a range of vertical and lateral axon– coil distances. Rows: results for the monophasic and half-sine stimulation waveforms, grouped by the three coil configurations. For the aligned figure-8 coil, the left region (negative x) is not fully shown due to the symmetry of the threshold distributions. Left and center columns: threshold values obtained with the conventional and modified cable equations, respectively. Right column: threshold difference comparing modified and conventional cable equations; any unmarked contour lines are spaced 10% and 5% apart for monophasic and half-sine waveforms, respectively. The outlines of the coils are illustrated as gray boxes, with the idealized windings located in the horizontal plane of y = 0. Color scales are the same within each column and shown at the bottom.

Polarization of unmyelinated axons by the longitudinal and transverse field components (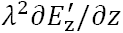 and 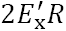) had similar amplitudes, given the field distribution presented in Fig. 2 and *λ*^2^/*R = r*_m_*σ*_i_/2 on the order of 10 mm for typical neuronal parameters. The modified CE (center column) indeed showed substantially different threshold distributions compared to the conventional CE; for a given vertical distance, stimulation thresholds at lateral positions with strong transverse field strength (i.e., under the center of the circular windings) were lower than thresholds at locations with large activating function (under the edge of the windings). Thresholds for the modified CE were substantially reduced compared to the conventional CE (right column), except when the axon was placed close to the center (within 1 cm lateral distance) of the aligned figure-8 coil.

The threshold distributions were qualitatively similar with the two waveforms, but quantitative changes were opposite for the conventional and modified CEs. Thresholds obtained with the conventional CE were higher for the half-sine waveform than the monophasic waveform due to the counteraction of the second phase on membrane polarization. For the modified CE, however, thresholds for the half-sine waveform were lower as a results of the transverse-field activation (excluding axon locations close to the center of the aligned figure-8 coil). Therefore, threshold changes due to transverse polarization were even more prominent for the half-sine waveform.

### Transverse stimulation of myelinated axons

The activation thresholds of straight myelinated model axons (Fig. 4) were substantially lower than those for the HH model and readily achievable with conventional magnetic stimulation devices. In myelinated axons, the longitudinal polarization was much stronger, with *λ*^2^ scaled approximately by the ratio of internodal and nodal length compared to unmyelinated axons (65). On the other hand, transverse polarization was present only at the nodes, and did not affect thresholds except when the axon was placed within a few millimeters of the coil center for the circular and perpendicular figure-8 coils. For the half-sine waveform, thresholds distributions were similar to those of the monophasic waveform, and the overall increase in threshold amplitude was in agreement with stimulation by the activating function. A qualitative difference occurred only for the perpendicular-orientated figure-8 coil, for which the half-sine waveform had a more symmetric distribution in the lateral direction (i.e., thresholds for coil positions in negative x-direction decreased to the same value as for positive positions).

**FIGURE 4.**
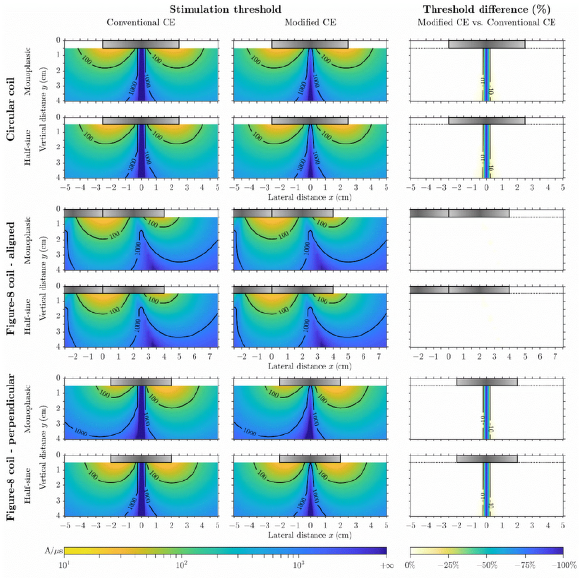
Activation thresholds for straight myelinated RMG-model axon. Similar format as Fig. 3, with a different color scale range for thresholds.

### Effect of axon undulation on activation threshold

The threshold distributions of the undulating axon were more uniform compared to those of straight axons, due the mixture of orientations of the axonal segments (Fig. 5, row 1 to 3). The axon undulation, by itself or with fascicle undulation present, reduced thresholds for nerves located laterally near the center of the circular windings (Fig. 5, row 4 and 6). The strong 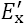 increased longitudinal activation (comparing 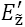 versus 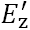), and “transverse-field activation” by 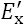 alone occurred when the nerve was perfectly aligned through the center of the circular and perpendicular figure-8 coils. Elsewhere, axon undulation increased thresholds because the projection of 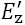 along the undulating path and weak 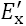 resulted in smaller 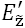 and decreased longitudinal activation. Fascicle undulation alone had weaker modulation effects on thresholds and mostly reduced thresholds under the center of coil windings (Fig. 5, row 5). For the half-sine waveform, the threshold distributions were qualitatively similar to those for monophasic stimulation, however, threshold modulation was weaker and fascicle undulation only reduced thresholds (figure available online).

**FIGURE 5.**
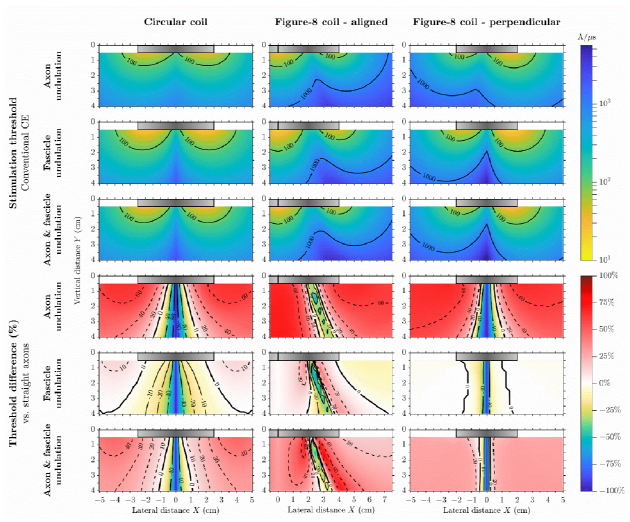
Activation thresholds of undulating myelinated RMG model axon for the three coil configurations (columns) using monophasic waveform. First, second, and third rows: threshold values for axons having only axon undulation, only fascicle undulation, and both undulation components. Fourth, fifth, and sixth rows: threshold difference compared to straight axons (versus left column of Fig. 4) for the three undulating axon models in the first, second and third rows; contour lines for positive and zero difference are shown with dashed and thick solid lines, respectively. Color scales are the same within each row and shown on the right; the same colors as in Fig. 4 are used for the thresholds and negative threshold differences.

## DISCUSSION

### Transverse polarization and electromagnetic coupling to neuronal membranes

As in electrical stimulation, both the transverse and longitudinal components of a magnetically-induced E-field can couple to neurons. Transverse polarization occurs on a sub-μs time scale and determines the deviation from the average membrane potential at a given longitudinal location of long neural cables. Based on the analysis of transverse polarization, the modified CE incorporates the effects of both E-field components and defines the activating function according to the relationship of the transversely-averaged potentials in the longitudinal direction (37). By applying these transverse and longitudinal relationships to E-fields induced by time-varying magnetic fields, we provided a rigorous theoretical justification for the CE for magnetic stimulation, including the use of scalar potentials for the transmembrane quantities, the inclusion of the transverse E-field, the substitution of the intracellular vector potential by the extracellular vector potential, as well as the use of quasi-potentials to drive the CE.

### Quasi-potentials for simulation of magnetic stimulation

We defined and theoretically justified the concept of quasi-potentials which allow simple and practical computational implementation of the CE for magnetic stimulation, for example by using built-in CE solvers such as the `extracellular` mechanism in NEURON (66). An alternative method for applying an exogenous E-field to neural cable models is to convert the activating function into equivalent intracellular currents applied to each neuronal compartment (13, 41, 42, 64). However, instead of the continuous first-order partial differential of the E-field, the activating function in computational models needs to be discretized as a first-order difference with asymmetric weights (i.e., intra-compartmental conductance) along the cable due to variation in neuronal parameters (13, 64, 67), modified at terminals to reflect a sealed boundary condition (13, 41, 67–69), and assigned additional terms at branching points governed by the current balance equation (13, 41, 69, 70). Neglecting these considerations (35, 71) results in “leaky” compartments in which the longitudinal current driven by the induced E-field may arbitrarily enter or exit without crossing the membrane and may therefore lead to significant discrepancies in the interpretation and conclusions regarding activation of neurons by magnetic stimulation. Thus, the current injection approach is more complex and prone to implementation errors than the quasi-potential approach, which does not require information about the neuron model properties other than the locations of the compartments and their topological connection.

### Transverse-field activation of peripheral nerves

The relative threshold distribution obtained with the modified CE for unmyelinated axons were seemingly in agreement with the experimental observation by Ruohonen et al. and their exploration using a modified activating function (44). However, the amplitudes required to activate unmyelinated axons were orders of magnitude above experimentally observed values. Indeed, as responses were characterized with electromyograms evoked from the median nerve (44), they were not due to activation of unmyelinated fibers, but rather large diameter myelinated Aα axons. Similarly, *in vitro* electrical recordings of compound action potentials from phrenic nerves are unlikely to reflect contributions from unmyelinated axons (48, 49). Therefore, transverse-field activation of unmyelinated axons cannot explain the experimental results, and the use of unmyelinated axons in magnetic stimulation models (23, 24, 29, 35) should generally be avoided unless specifically modeling such axons or diseased nerves with demyelination.

In contrast to the unmyelinated axon model, the myelinated axon model indicated that transverse stimulation did not affect thresholds in straight myelinated axons and the threshold was always lowest for coil positions where the conventional activating function was largest. Therefore, the results from both membrane models indicated that the polarization due to transverse E-field alone cannot account for the experimental observations.

Apparent transverse-field stimulation is likely the result of an E-field component along the axon that generates an activating function due to factors such as the geometry of the tissue surrounding and within the nerve, electrical inhomogeneity and anisotropy, and variations in the geometrical and/or electrical properties of the axon. For instance, the undulations of axons and fascicles result in short and curved axonal segments, locally aligned with E-field components that are globally transverse to the nerve (49, 54). The effect of undulation on thresholds was stronger for axon undulation, but fascicle undulation contributed as well. Undulations in the nerve reduced the thresholds for coil positions where the activating function longitudinal to the overall nerve trunk orientation was small, thereby creating apparent transverse field stimulation. Further, nerve undulation increased thresholds for coil positions where the longitudinal activating function was large. Therefore, nerve undulation flattened the threshold distributions for all coil configurations and reduced the sensitivity of threshold to coil position. For a fixed vertical axon–coil distance, the differential effect of undulation on the thresholds for various lateral coil positions resulted in threshold profiles resembling those recorded experimentally (44).

### Limitations

The simulation of magnetic stimulation in this paper had several simplifying assumptions, such as the use of semi-infinite volume conductor with homogenous conductivity (44). The E-field is influenced by the secondary charges on the boundary of the volume conductor and internal discontinuities. For example, both the cylindrical shape of the forearm and the perineurium of individual fascicles shape the E-field distribution. The cylindrical boundaries reduce the field amplitude but have less effect on the distribution (44, 72). Considering the attenuation of the E-field amplitude due to the perineurium, especially in the transverse direction, the contribution of the transverse field to activation of straight axons would be further reduced. Therefore, the conclusion that transverse polarization alone is insufficient for transverse stimulation remains valid even with these simplifications.

The reduced transverse field also affects the E-field in undulating axons but to a lesser extent. The undulation of axons is not confined to one plane, and can, for example, exhibit helical shapes that manifest as bands of Fontana (55, 56). As axons leave and join the nerve or the number and arrangement of fascicles changes over distance, the undulation of an individual axon is likely more irregular, not having steady sinusoidal variations with fixed amplitudes and wavelengths. The depth of the median nerve varies within the forearm, and the variation of the distance to the coil could further affect stimulation thresholds. The undulation of the axon was the major contributor to neural activation by the E-field transverse to the nerve trunk. However, the parameters for the undulation were drawn from previous histological measurements that could deviate from the actual geometry due to tissue deformation. Especially, shrinkage of the nerve could result in underestimates of the amplitude and overestimates of the wavelength of the undulation, which would, in turn, affect thresholds (58).

## CONCLUSIONS

Transverse polarization and the modified CE provided a sound theoretical foundation and an improved theoretical justification for the use of the CE and quasi-potentials in simulations of magnetic stimulation. Using this theoretical framework, the effects of transverse polarization and nerve undulation on activation threshold were studied. While the thresholds for unmyelinated axons were affected by transverse polarization, this was not the case for unmyelinated axons. Therefore, the experimentally observed activation of nerves by transverse E-fields cannot be explained by transverse axonal polarization but is likely due to nerve fiber undulation and other spatial inhomogeneities that cause a local longitudinal E-field in the axon.

## AUTHOR CONTRIBUTIONS

A.V.P. and W.M.G. conceived and supervised the study. B.W. performed the theoretical analysis, computational models, and simulations; analyzed data; and generated figures. B.W. wrote the manuscript; A.V.P. and W.M.G. revised the manuscript; all authors approved the final submitted version.

## ACKNOWLEDGMENT

This work was supported by Grant No. R01NS088674 from the National Institutes of Health and a Research Grant by Tal Medical. The simulations in this work utilized the Duke Compute Cluster. The authors thank Aman S. Aberra and Dr. Stefan M. Goetz for discussion of electromagnetic–neuronal coupling.

Preliminary results of this study were presented at the 6th International Conference on Transcranial Brain Stimulation (September 2016, Göttingen, Germany) (73) and at the 47th Annual Meeting of the Society for Neuroscience (November 2017, Washington DC, United States of America) (74).

B.W. is inventor on patent applications on technology for transcranial magnetic stimulation. W.M.G. has no relevant conflicts to report. A.V.P. is inventor on patents and patent applications, and has received research and travel support as well as patent royalties from Rogue Research; research and travel support, consulting fees, as well as equipment loan from Tal Medical; patent application and research support from Magstim; and equipment loans from MagVenture, all related to technology for transcranial magnetic stimulation.

## APPENDIX A CHARGE STORAGE ON SHEET OF CELL MEMBRANE

A planar sheet of membrane in the vertical plane is considered, with the intra-and extra-cellular spaces towards the left and right, respectively (Fig. A1). Only the transverse component of the E-field is considered, which is normal to the surface and has strength of *E*′. The conductivity and permittivity are *σ*_i,e_ and *ε*_i,e_, with the subscripts i and e indicating intra-and extracellular, respectively. The membrane has specific capacitance *c*_m_ = *ε*_m_/*d* and specific resistance *r*_m_ *= d*/*σ*_m_, where the *σ*_m_ and *ε*_m_ are the intramembrane conductivity and permittivity, respectively, and *d* is the membrane thickness that is negligible compared to the neuronal size.

Due to the difference in conductivities in intra-and extracellular spaces, the current densities driven by the primary field are not equal in the two domains, resulting in a net charge density *q*_s_ stored on the membrane. The time scale of this process *τ*_s_ is a combination of the time constants of the intra-and extra-cellular spaces *τ*_i,e_ = *ε*_i,e_/*σ*_i,e_ and they are all on the order of tens to hundreds of picoseconds. Therefore, charge storage is considered instantaneous and continuity of normal current density on the boundary of the two domains (19, 29) is ensured by the secondary E-field 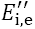 (orientation as defined in Fig. A1)

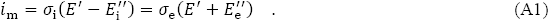

According to Gauss’s law, the stored charge density and the E-fields satisfy the relationship

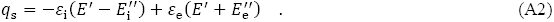

The exact solution for the three quantities 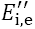 of the solution space and the distribution of the primary E-field. However, due to the extremely low conductivity of the membrane (*σ*_m_ ≪ *σ*_i,e_), the charge carriers in the intra-and extracellular spaces do not simply enter the other side during the charge storage process. Therefore, the net charge storage consists of charge *q*_i_ and *q*_e_ of opposite polarity accumulated on the intra-and extracellular sides

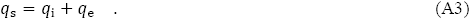

The differential of the single-sided charge density

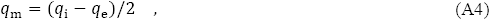

establishes a secondary intramembrane E-field 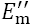 across the membrane without affecting the E-field in the intra-and extracellular spaces

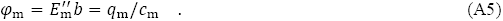

The transmembrane potential *φ*_m_ is given without the double prime for simplicity, because the primary field, regardless of origin is considered continuous.

The behavior of the membrane polarization (*q*_m_ and *φ*_m_) is described by a differential equation on the time scale of *τ*_m_, for which the instantaneous solutions to Eqs. A1–A5 provide the initial state *φ*_m,0_ and the input *i*_m_. Thus, the membrane polarization can be decomposed into two independent components. The first component is the zero input response *φ*_m,zi_, which dissipates *q*_m,0_ through the membrane to zero so that the single-sided charge density reaches an equilibrium of *q*_i,e_ *= q*_s_/2 and then reverses polarity and cancels the primary field in the membrane. The amplitude of *φ*_m,zi_ throughout this entire process is on the order of nanovolts per unit applied field. Therefore, this component is negligible compared to typical membrane responses to extracellular stimulation, whose amplitude is on the order of micro-to millivolts per unit applied field (75). The second membrane polarization component

*φ*_m_ = *φ*_M,zs_ starts with no polarization and is the zero state response. This component is determined by the secondary E-field in the membrane only and is the response to the current density input Eq. 3 determined by the current continuity Eq. A1 with a steady state of 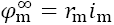 For typical polarization of millivolts, the secondary E-field associated with this component is order of magnitudes larger than the applied fields (∼ 10^7^ V/m versus < 10^3^ V/m).

This analysis shows that the membrane polarization contains a very small component *φ*_m,zi_ that is a transient response due to charge storage and is not part of the current continuity equation. Because the source field in electrical stimulation and the secondary field by the membrane can both be represented by scalar potentials, they can be simply added to cancel out (see right panel of Fig. A1). However, the source field of magnetic stimulation contains a non-conservative component that complicates the theoretical consideration for the transmembrane potential, which can be addressed using quasi-potentials as discussed in Methods section.

## APPENDIX B PARAMETERS OF NONLINEAR NEURONAL MEMBRANE MODELS

The HH model (63) was adjusted to room temperature (23.5°C) with *Q*_10_ = 3, and a radius of 3 μm was used following previous studies (37, 38, 53), with longitudinal discretization set to 82.1 μm according to the d_lambda (d*λ*) rule of NEURON (66) with d*λ* = 0.1. The RMG model (64), based on human peripheral nerve fibers at body temperature (37°C), had fast sodium channels, persistent sodium channels, and slow potassium channels at nodes, which are each modeled as a single compartment. The passive internodes had the same radius as the unmyelinated axon and corresponded to a 10 μm outer fiber diameter including the myelin; the membrane properties were calculated for 120 myelin lamella and the internodes were discretized into ten and hundred compartments for straight and undulating axons, respectively. The axon geometry and passive electrical parameters for the two models are summarized in Table B1. For the active ion channel parameters and dynamics, please refer to the original publications of the respective membrane models (63, 64). Detailed information can also be found in the code available online.

**FIGURE A1.**
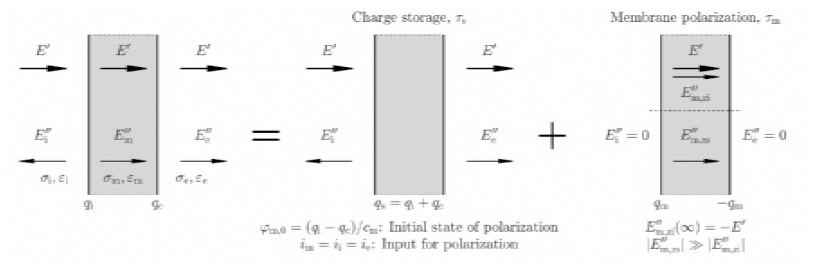
Illustration of two independent behaviors of the cell membrane in response to externally applied (primary) E-field. The secondary field generated in response to the primary field can be decomposed into charge storage (middle) and membrane polarization (right). The depicted arrows are not proportional to the E-field strength and only illustrate direction. The charge densities involved in each case are given at the bottom. The charge storage occurs on a very fast time scale, and results in current continuity across the membrane, providing the input for membrane polarization. On the other hand, the charge is not equal on the two sides due to the extremely resistive membrane and establishes a temporary polarization that becomes the initial state for the membrane. The membrane polarization therefore can be considered as two components, the zero input response (above the dashed line) that dissipates the initial state and also cancels the primary field in the membrane, and the zero state response (below the dashed line) which polarizes the membrane according to the current continuity equation.

**TABLE B1.**
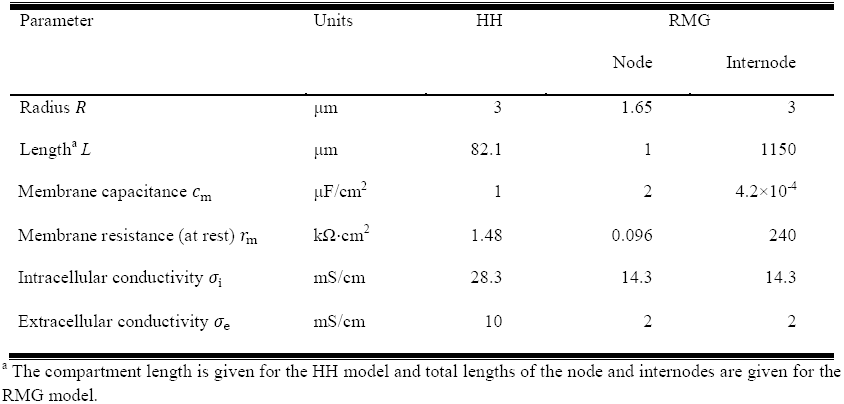
Axon parameters of unmyelinated (HH) and myelinated (RMG) axons.

